# A semi-automated pipeline for morphological analysis of myonuclei along single muscle fibers

**DOI:** 10.64898/2025.12.15.694446

**Authors:** Esben T. Schroeder, Helia G. Megowan, Madeline Luu, Adam Shuaib, Adam C. Fries, Jake Searcy, Hans C. Dreyer

## Abstract

Manual quantitation of skeletal muscle myonuclear number, spatial orientation, and morphology is time-consuming and subject to error and bias. To overcome these limitations, we developed and validated a semi-automated, quantitative, and reproducible image-analysis pipeline. The workflow combines FIJI-based preprocessing with custom Python scripts to process immunohistological images of individual muscle fibers, enabling high-resolution and scalable quantification of nuclei. Analyses incorporate morphometric parameters including nuclear position, shape, and three-dimensional orientation, as well as centroid-to-skeleton distance and nearest-neighbor relationships to capture spatial patterns of myonuclear organization along the fiber. Outputs include per-fiber and biopsy-level summaries integrated with *Imaris* metrics. This semi-automated approach provides a robust and efficient platform for high-throughput analysis of myonuclear number and structural features across large single fiber datasets.

## Introduction

Histological analysis of skeletal muscle tissue is helpful to understanding the relationship between anatomical structure and physiological function^1^. Immunohistochemical techniques, which employ labeled antibodies to localize specific proteins, are often integrated with advanced imaging methods to provide detailed insight into the structural organization of muscle tissue. Muscle fibers exhibit remarkable plasticity in response to internal and external stimuli, leading to morphological adaptations such as increases in muscle fiber cross-sectional area (CSA), changes in mitochondrial and myonuclear content, and fiber type transitions. These structural adaptations visualized histologically are commonly used as proxies for functional capacity.

A prevailing method in the muscle field is to quantify structural and positional changes histologically using tissue sections cut perpendicular to the long axis of the fibers. This cross-sectional approach allows for more individual cells/fibers (each fiber is a single muscle cell) to be analyzed providing greater coverage of the muscle being studied and therefore better representation. However, if information about myonuclei number and structure are of interest, the cross-sectional approach may under sample given the multinucleated syncytial nature of muscle cells. For example, along a one millimeter of muscle cell there are 100 10-µm thick cross sections and if the muscle bundle obtained from biopsy are 3 cm to 5 cm in length^2,3^, this constitutes 3000 to 5000 10-µm thick samples. Said another way, one 10-µm thick cross section represents 0.00033% (1/3000) to 0.0002% (1/5000) of the approximate 3-5 cm length of single muscle fiber cells^2,3^—and averages out to about 2 to 3 myonuclei per each 7-µm to 10-µm thick cross section. Conversely, hundreds to thousands of myonuclei can be included when sampling lengthwise along the long axis of single muscle fibers, improving statistical power, confidence, and rigor for studies that center on the changes to myonuclear structure and function.

Precise quantification of myonuclei is critical for advancing our understanding of skeletal muscle biology, particularly in relation to adaptations such as myonuclear accretion. This is especially relevant to the ongoing assessment of the myonuclear domain hypothesis, which posits that each myonuclei governs a defined cytoplasmic volume^4,5^. Traditionally, myonuclear domain has been assessed using cross-sectional tissue samples; however, recent studies suggest that myonuclear accretion correlates more strongly with myofiber surface area rather than fiber volume alone^6–8^. Beyond myonuclear accretion, additional parameters that may be of interest to the muscle biology field—including myonuclear shape, spatial orientation along the fiber’s long axis, and the distribution and/or clustering of myonuclei of similar shape—are emerging as important indicators of muscle cell regulation and remodeling^9–12^.

Motivated by the above, in this study, we developed a semi-automated image analysis pipeline for immunohistological images of isolated single muscle fibers that allows:

1. *Quantification of myonuclear number along the length of individual fibers*,
2. *Classification of each myonucleus into predefined morphological categories*,
3. *Assessment of local clustering by quantifying the number of similarly shaped myonuclei within a fixed radius*, and
4. *Quantification of the long-axis orientation of each myonucleus relative to the long-axis of the single fiber*.

To our knowledge, this is the first study to integrate these analytical features into a single pipeline using widefield microscopy—a platform broadly accessible in core facilities and individual laboratories. This approach allows high-throughput analysis of large composite images, yielding rich datasets on nuclear organization and spatial relationships along individual myofibers. The method has broad applicability in biomedical research and may help uncover myonuclear structure-function relationships within various physiological and pathological contexts across the lifespan.

## Methods and Results

### Ethical approval

The study was approved by the University of Oregon Institutional Review Board and conducted in accordance with *Declaration of Helsinki* ethical principles. All subjects gave informed written consent prior to study participation.

### Biopsies

Skeletal muscle biopsies were performed on human vastus lateralis in a distal to proximal sequence (if multiple biopsies) and in the fasted state. For serial biopsies the biopsy needle was oriented lateral from and perpendicular to the long axis of the femur with the biopsy needle window/opening situated in the mid-substance of the muscle tissue. Serial biopsies were thus obtained from the mid-substance of the vastus lateralis muscle that are 3 or more cm apart (and not angled)^13,14^. Biopsy samples were obtained using a Bergström needle with 5mm internal diameter with manual suction. Each biopsy yielded approximately 300 mg of tissue (range ∼100—500 mg) with the fiber bundles approximating 2 to 7 mm in length. Our goal for single fiber analysis is to obtain a minimum of 20 individual muscle cells for analysis per biopsy.

### Single fiber isolation and immunohistochemistry

Immediately following biopsy collection, tissue sections with the longest fiber bundles were teased apart and placed in a 0.6 ml Eppendorf tube containing 500 µl 4% paraformaldehyde (PFA) for 48 hours at 4ºC. Fixed fiber bundles were then placed in a 0.6 ml Eppendorf tube containing PBS and kept in the dark at 4ºC until further processing.

To begin tissue processing, the biopsy sample was placed into a 3 cm diameter glass dish containing a small amount of PBS to prevent drying out. Under a benchtop stereomicroscope, the fixed sample was teased apart with forceps into smaller bundles (final size being ≤ 30 fibers), with care being taken to avoid damaging or tearing the fibers. Once separated, the next step is to remove all extracellular components from the sample to further isolate the single fibers. To achieve this, the tissue bundles were transferred into a 1.5 ml Eppendorf tube containing 750 µl of 40% NaOH, then placed on a tube rotator at moderate speed for 2 hours to facilitate digestion of the connective tissue between the fibers. Following digestion, NaOH was removed by filtering the bundle through a 40 µm mesh strainer and repeatedly washed with 1x PBS (10-12 total). Fibers were then collected from the strainer using forceps and transferred into a 0.6 ml Eppendorf tube containing 300 µl 1x PBS until further processing.

Following isolation, fibers were permeabilized in a 1.5 ml Eppendorf tube containing 750 µl 1% Triton-X100 for 10 minutes on the rotor wheel. Permeabilized fibers were then centrifuged at 13,000g for 3 minutes to pellet fibers. After centrifugation, supernatant was removed, and fibers were incubated in primary antibody mouse anti-slow myosin (cat. no. M8421, Sigma-Aldrich, Darmstadt, Germany), 1:200 in PBS for 90 minutes at room temperature. Following incubation in the primary antibody, fibers were washed three times for three minutes in 1% Triton-X100, centrifuged (13,000g for 3 minutes to pellet fibers). Fibers were then transferred to a 1.5 ml Eppendorf tube containing the secondary antibody Alexa Fluor 555 goat anti-mouse IgG1 (cat. no. A-21127, Invitrogen, Carlsbad, CA, USA), 2:500 in 250 µl PBS and 250 µl 1% Triton-X100 for 1 hour at room temperature. Following incubation in the secondary antibody, fibers were again washed three times for three minutes in 1% Triton-X100, with centrifugation. Fibers were then placed in 100 µl of PBS with 1 µl of 100 µM DAPI (DAPI Hydrochloride, cat. no. D1306, Invitrogen, Carlsbad, CA, USA) for 5 minutes.

Following staining, fibers were transferred to glass microscope slides and carefully separated with forceps to avoid overlap. Vectashield Mounting Medium (H-1000, Vector Laboratories, Newark, CA, USA) was applied to maintain separation and prevent fading. Slides were then mounted with a glass coverslip and sealed with clear nail polish.

### Image acquisition

All slides of fluorescently labeled, fixed single fibers were imaged using a Leica DMi8 Thunder wide field fluorescence microscope equipped with Leica’s LED8 illuminator and K8 camera (6.5 µm pixels). Fibers were first visually characterized as either Type I (MyHC-1^+^) or Type II (MyHC-1^-^) using the HC PL FLUOTAR 10x/.32 objective in the TRITC channel (excitation and emission filters: 554/24, 594/32) at 25% illumination intensity and an exposure time of 200 ms (Figure 1).

**Figure 1:**
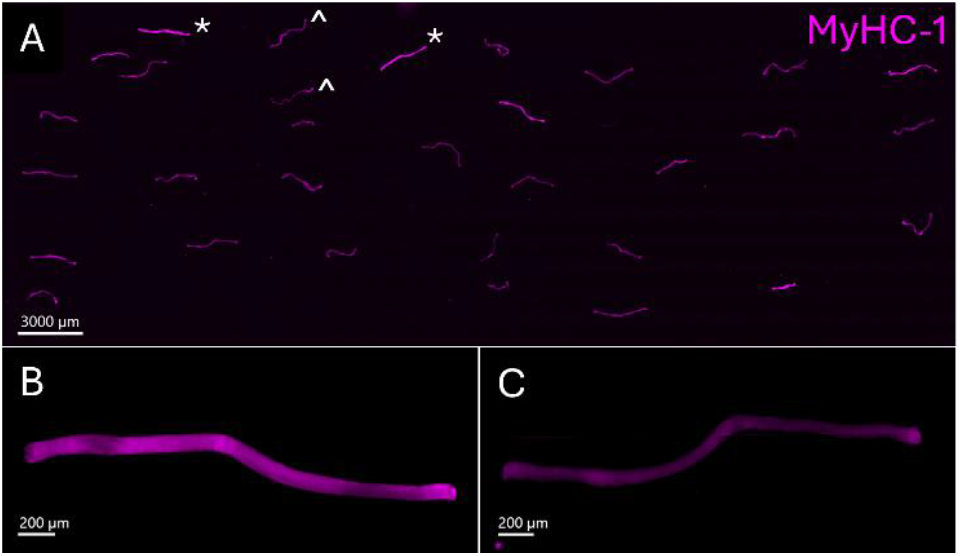
Fiber Type Characterization. **(A)** Representative merged Tile Scan image of fiber type characterization for a slide of fixed muscle fibers stained with Myosin Heavy Chain I (MyHC-1); fibers are characterized based on fluorescence intensity under the TRITC channel (555 nm). Type I fibers (*) fluoresce at a greater intensity than Type II fibers (^), scale bar = 3000 µm. **(B)** Representative type I muscle fiber. **(C)** Representative type II muscle fiber.

Once identified based on fiber type, image stacks of myonuclei on entire single fibers spanning multiple fields were acquired at higher magnification and resolution using a HC PL FLUOTAR 20x/0.4 objective in the DAPI channel (excitation and emission filters: 407/375, 450/20) at 27% illumination intensity, exposure time of 100 ms, and 2.7 µm step size (SVI Nyquist calc., https://svi.nl/Nyquist-Calculator). Image stacks were then stitched in the LAS X software for downstream analysis.

### 3D myonuclear quantification using Imaris

Raw, merged z-stacks of single fibers (Figure 2A) were first converted to the.*ims* format before being imported into *Imaris* for further analysis. Prior to myonuclear quantification, images were processed with a background subtraction algorithm (filter width = 10 µm) to enhance myonuclear signal relative to the fiber background (Figure 2B). Myonuclei from multiple single fibers were measured using a Line Profile within *Imaris*, from which the average diameter of a myonucleus was estimated to be 10 µm, resulting in a background subtraction filter width of 10 µm.

**Figure 2:**
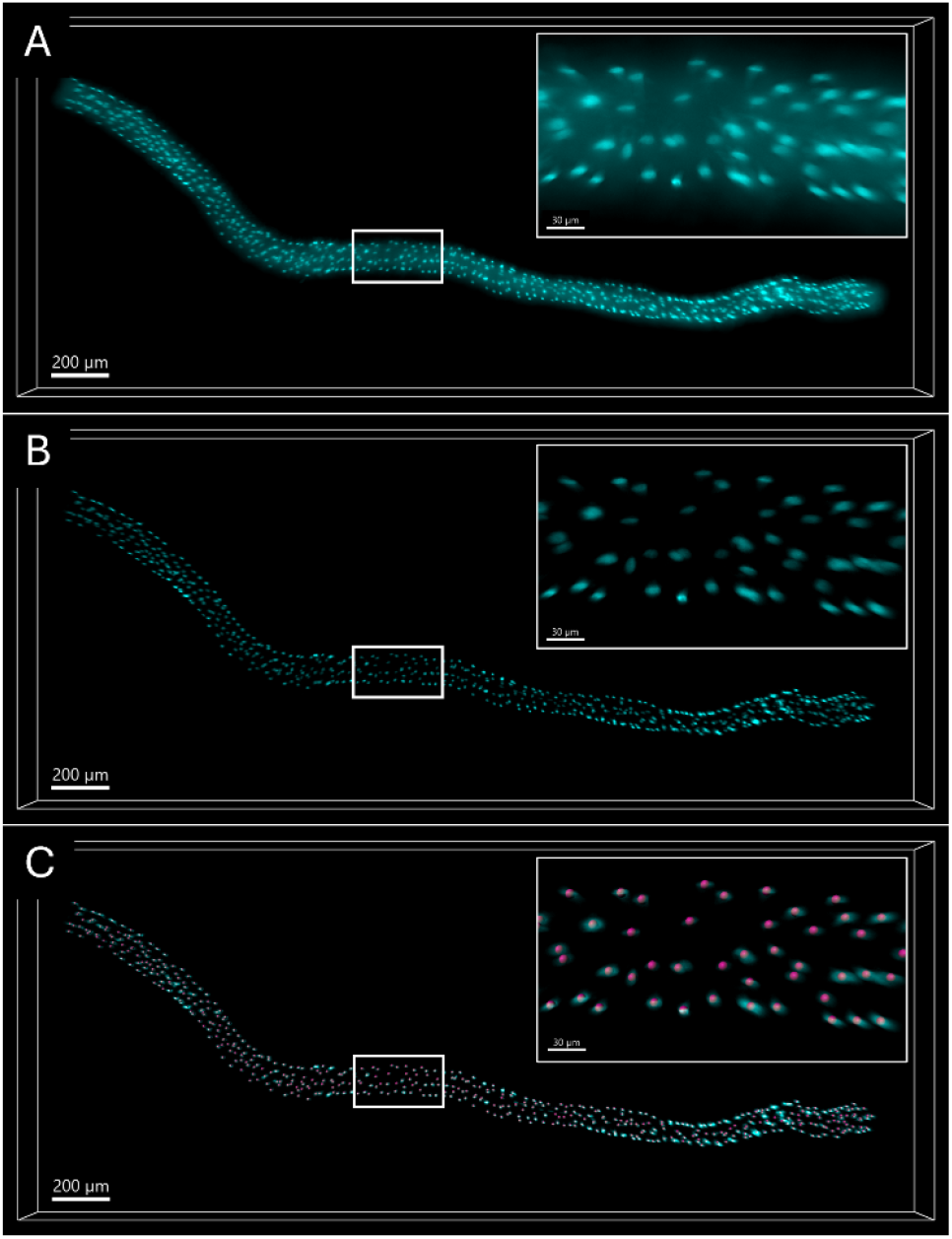
Myonuclear Quantification in Imaris. **(A)** Raw z-stack of a single muscle fiber acquired under the DAPI channel (cyan). **(B)** Background-subtracted DAPI channel (filter width = 10 µm). **(C)** Myonuclear quantification on single muscle fiber; *Spot* (magenta) placed at center of each identified myonucleus (cyan).

*Imaris’ Spots* module was used for accurate myonuclear quantification. First, the *Estimated XY Diameter* of the feature object was input as 10 µm based on previously described myonuclear measurements. *Imaris’ Spots* prediction was then thresholded using *Imaris’ Quality* parameter to obtain myonuclear counts (Figure 2C). Total counts were manually quality-controlled by including counts of myonuclei not picked up by the *Spots* algorithm due to weak fluorescence signal, as well as manually excluding *Spots* that were incorrectly placed on aberrant DAPI signal or on nuclei outside of the muscle fiber.

When comparing *Imaris’* total myonuclear counts without manual quality control to counts with manual quality control on 622 fibers, a Bland-Altman analysis resulted in a bias of 3.821 and limits of agreement from -3.489 to 11.13 (Figure 3A). Across the 622 fibers, the average nuclei/mm of muscle tissue with manual correction was 171.9 compared to 165.3 without manual correction. The r value between methods was 0.9961 (Figure 3B) and the percent error was 4.02 × 10^-3^. *Imaris* tended to have lower counts without manual quality control, potentially due to nuclei clustering on top of each other within a single fiber that prevented *Spots* from accurately segmenting the clustered object as multiple individual myonuclei. As myonuclear number is a primary outcome for our projects, we chose to include this manual quality control.

**Figure 3:**
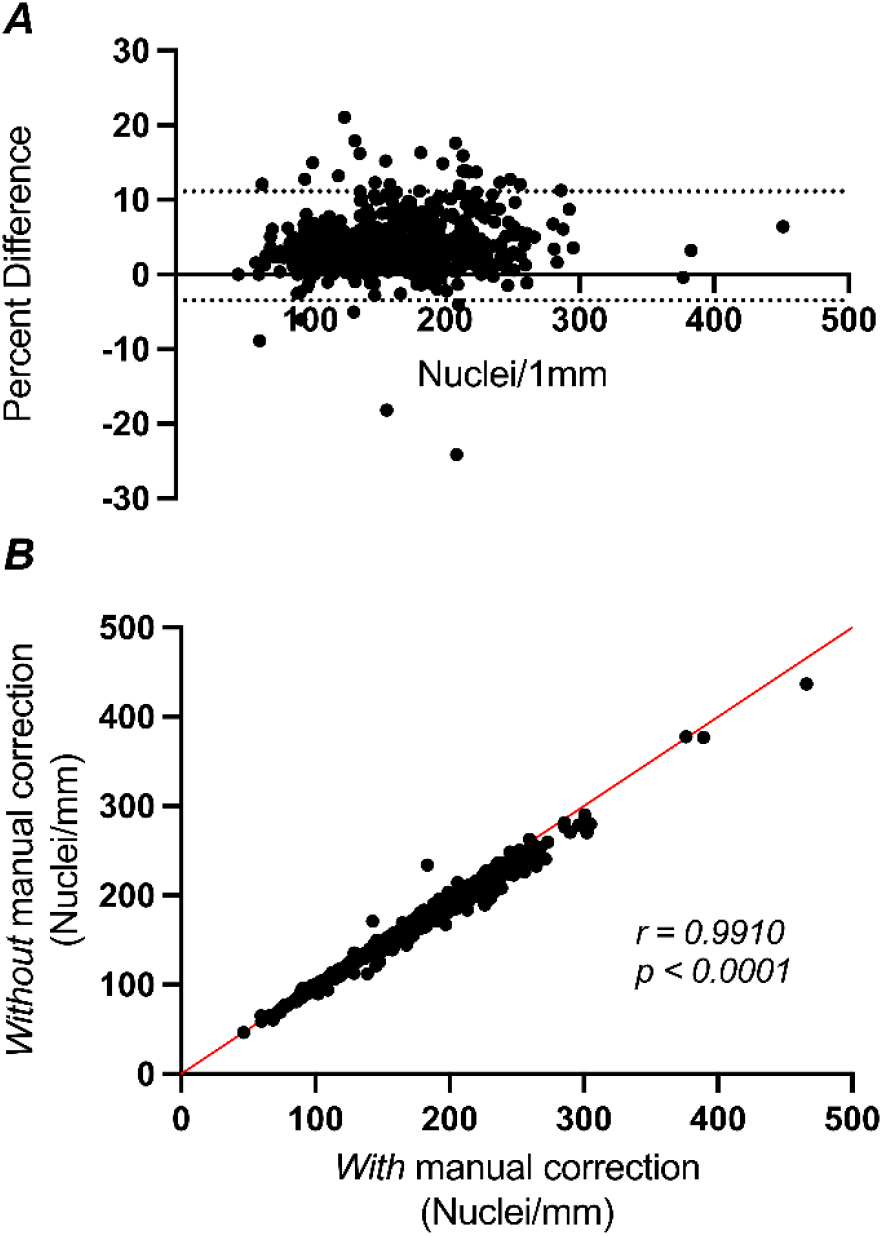
Comparison of nuclei counted per mm of muscle tissue from *Imaris* with and without manual quality control (N = 619). (**A**) Bland-Altman 95% limits of agreement plot comparing *Imaris’* nuclei counts with and without manual correction. (**B**) Correlation between nuclei / 1 mm of muscle tissue without and with manual correction; r = 0.9910 and p < 0.0001.

### 3D fiber length measurement

Fiber length is obtained using *Imaris’ Filaments* module. First, a mask of the entire fiber is created using the *Surface Creation* module. The whole single fiber is segmented on the DAPI channel using a smoothing width of 15 µm, then masked and binarized by setting the pixel value within the segmented surface to 1.00. These masking steps in the *Surface Creation* module result in a duplicated DAPI channel with consistent signal throughout the entire single muscle fiber in all 3 dimensions.

*Imaris* uses representative *Seed Points* placed along the fiber as a guide to estimate the path that the *Filament* should follow along the DAPI signal. This method optimizes both the source channel with a smoothed, binarized DAPI signal, as well as optimal *Seed Points* that are manually placed along the fiber. After indicating the *Thinnest Diameter* of the object of interest as 100 µm (approximate diameter of a muscle fiber), *Imaris’* default *Seed Points* are removed. Then, in *Slicer View* with a *Slice Width* of 10 µm, new *Seed Points* are manually placed at the center of the fiber from end-to-end. This results in a smoothed *Filament* (Figure 4) that follows the center of the muscle fiber. Fiber length is measured by obtaining the *Filament* length in microns.

**Figure 4:**
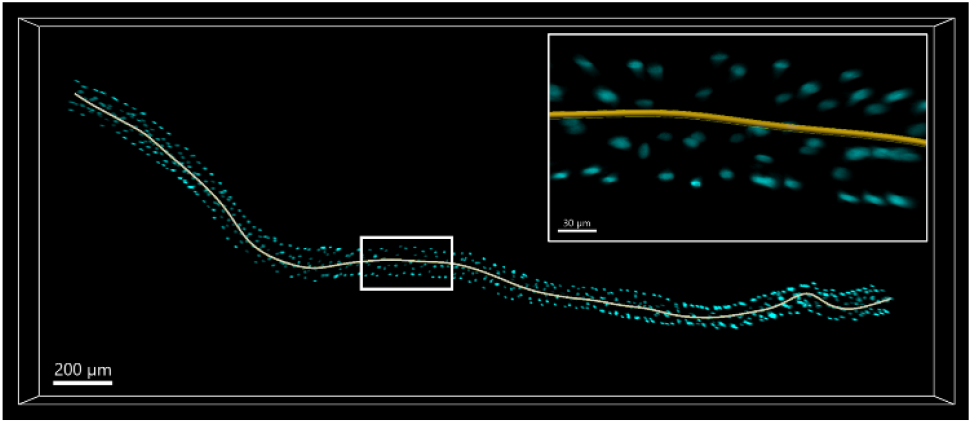
Fiber Length Measurement in *Imaris*. Single muscle fiber with a *Filament* (yellow) placed along the center of the fiber from end-to-end (fiber length = 3061 µm).

For validation of this method, 88 single muscle fibers were measured manually using 150 µm segments placed along the fiber’s central axis in *Imaris’* Slice View mode. Bland-Altman analysis between manual and *Imaris*-derived fiber lengths resulted in a bias of 0.279 and limits of agreement from -2.511 to 3.069 (Figure 5A). Across the 88 fibers tested, the average fiber length was 3054 µm when measured manually compared to 3045 µm when measured by *Imaris*. The r value between manual and *Imaris*-derived fiber lengths was 0.9997 (Figure 5B) and the percent error was 2.69 × 10^-5^. With such small differences between methods, we chose to go forward with the *Imaris*-derived fiber length measurements for our analyses.

**Figure 5:**
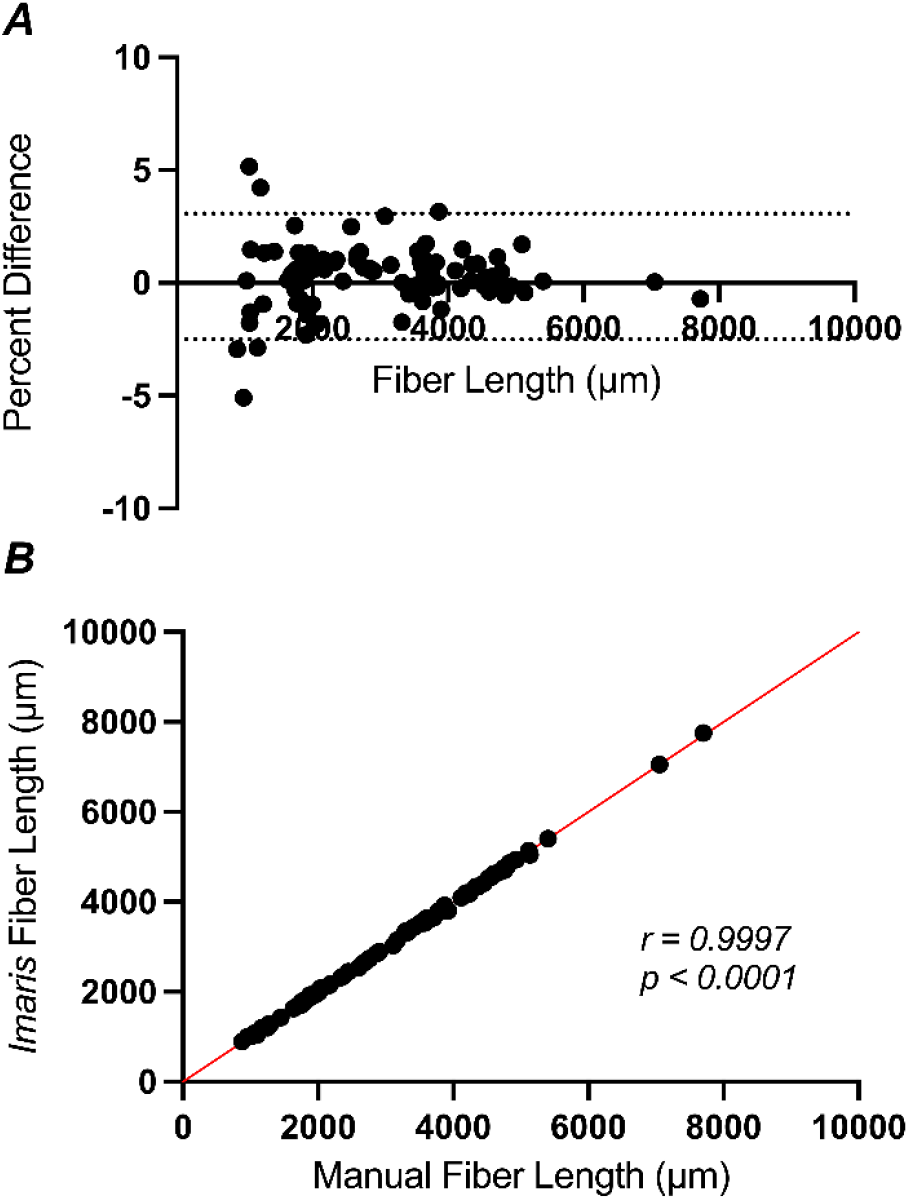
Comparison of fiber length in micrometers from *Imaris* and manual methods (N = 88). (**A**) Bland-Altman 95% limits of agreement plot comparing *Imaris’* fiber length measurement to manual measurement. (**B**) Correlation between fiber length from *Imaris* and manual; r = 0.9997 and p < 0.0001.

### 2D analysis of myonuclear shape, position, orientation, and co-localization

3D segmentation using 20x widefield z-stacked images in *Imaris* allowed us to efficiently quantify myonuclear density from a large number of fibers; however, when attempting to 3-dimensionally render myonuclear morphology from these images, z-axis elongation and excessive light scattering distorted the true shapes of the myonuclei. This distortion, an inherent issue with widefield systems, resulted in nuclei appearing artificially elongated in the z-dimension.

To glean information on myonuclear shape, position, and orientation, while still utilizing widefield images that can be collected with relative ease, we decided to pivot to a 2D analysis using 2D projections of the original 3D z-stacks. This 2D visualization allows us to meaningfully and accurately group myonuclei of similar shapes using aspect ratio – a metric that has been applied by other groups in the field^10,15–17^. Additionally, this approach allows us to capture information about myonuclear orientation and colocalization, all without the added artifacts that the z-dimension adds to these images.

To do this, we developed a fully automated macro pipeline in FIJI designed to convert each z-stack into a 2D standard deviation projection (STD projection) for efficient 2D morphometric analysis of each myonuclei. This approach collapses the z-stack into a single image that emphasizes variation in intensity across z-planes, effectively highlighting myonuclei while reducing the signal of unwanted artifacts produced in the z-dimension during image acquisition. Importantly, this also allows for robust thresholding and segmentation steps to be applied consistently across images, greatly improving throughput and reproducibility for efficient automated processing of large datasets.

### Overview of the FIJI macro pipeline

The macro was designed to handle batch processing of Leica.*lif* files and automatically parses sample metadata (SubjectID, timepoint, leg) to organize outputs. It runs fully automated in FIJI macro language batch mode and includes the following steps:

#### 1. Image Import and Preprocessing

Each.lif file is opened using Bio-Formats. The macro selects only those series labeled “Merged” for analysis. Prior to image processing, a raw TIFF of the z-stack (Figure 6A) is saved downstream z-plane identification of objects. A rolling ball background subtraction (radius = 30 pixels, the average diameter of a myonucleus) is applied to the z-stack to enhance nuclear contrast (Figure 6B).

**Figure 6:**
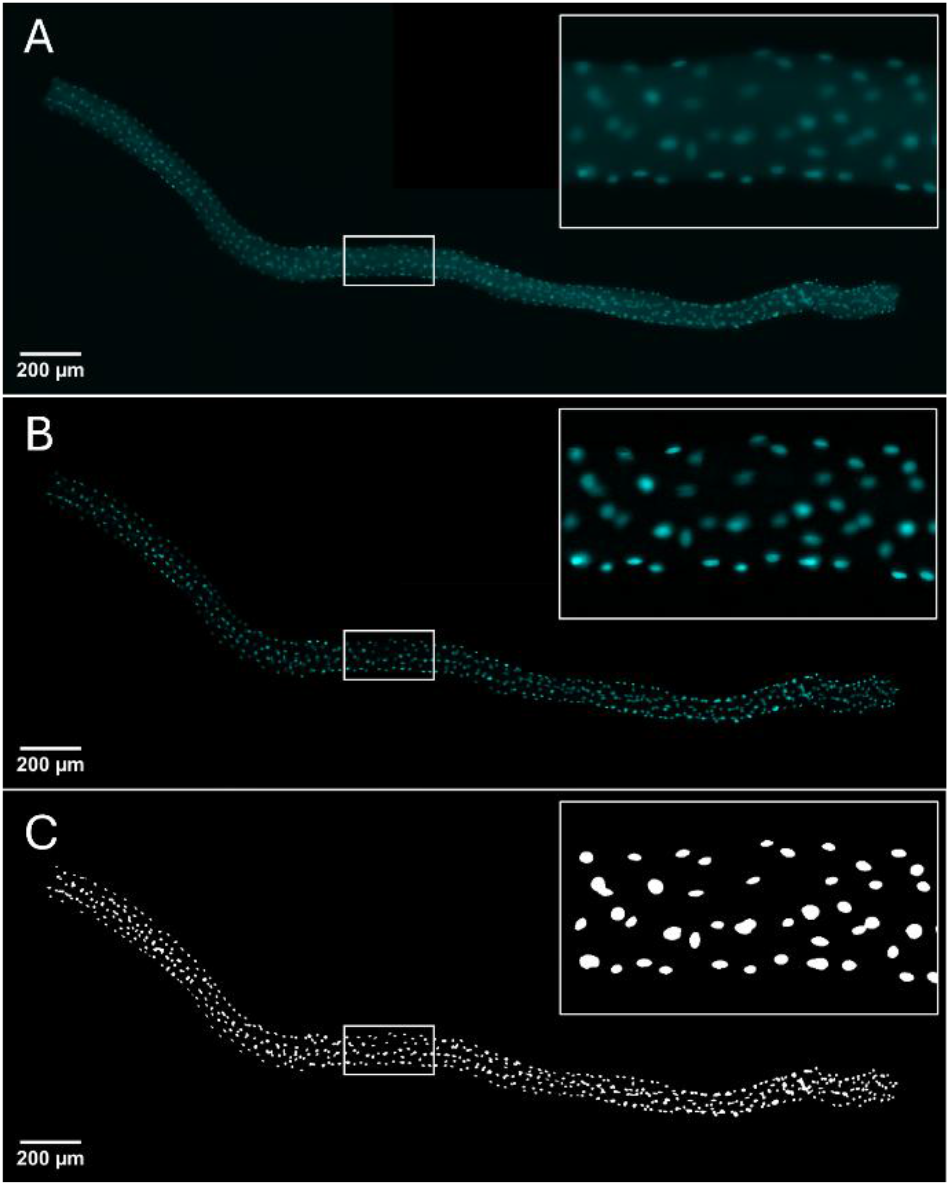
2D Nuclei Mask Generation. **(A)** Single muscle fiber z-stack acquired under the DAPI channel (358 nm). **(B)** Background-subtracted DAPI channel (filter width = 10 µm). **(C)** STD z-projection of the DAPI channel following binarization with Otsu thresholding + Median filter (filter width = 2 px).

#### 2. Standard Deviation Z-Projection

The macro generates a 2D projection using the “Standard Deviation” method, which accentuates objects that differ in mean intensity across z-slices, which is ideal for identifying myonuclei. A median filter (radius = 1 pixel) is applied to reduce noise before automatic thresholding via Otsu’s method. The image is binarized and converted to an 8-bit mask to ensure compatibility with downstream processing (Figure 6C). These projections are saved under a “STDIP” directory for each sample.

#### 3. Skeletonization Workflow

The projected image is duplicated, blurred (Figure 7A) and thresholded again with Otsu (Figure 7B). This step transforms the DAPI mask created in Step 2) into a representative mask of the overall fiber shape. FIJI’s *Analyze Particles* is then used to isolate objects larger than 70,000 pixels^2^, thus segmenting out any smaller objects remaining in the image. This mask is converted to 8-bit binary to guarantee compatibility with the Skeletonize function in FIJI (Figure 7C). The resulting 1-pixel-wide skeleton is saved under a “Skel” directory.

**Figure 7:**
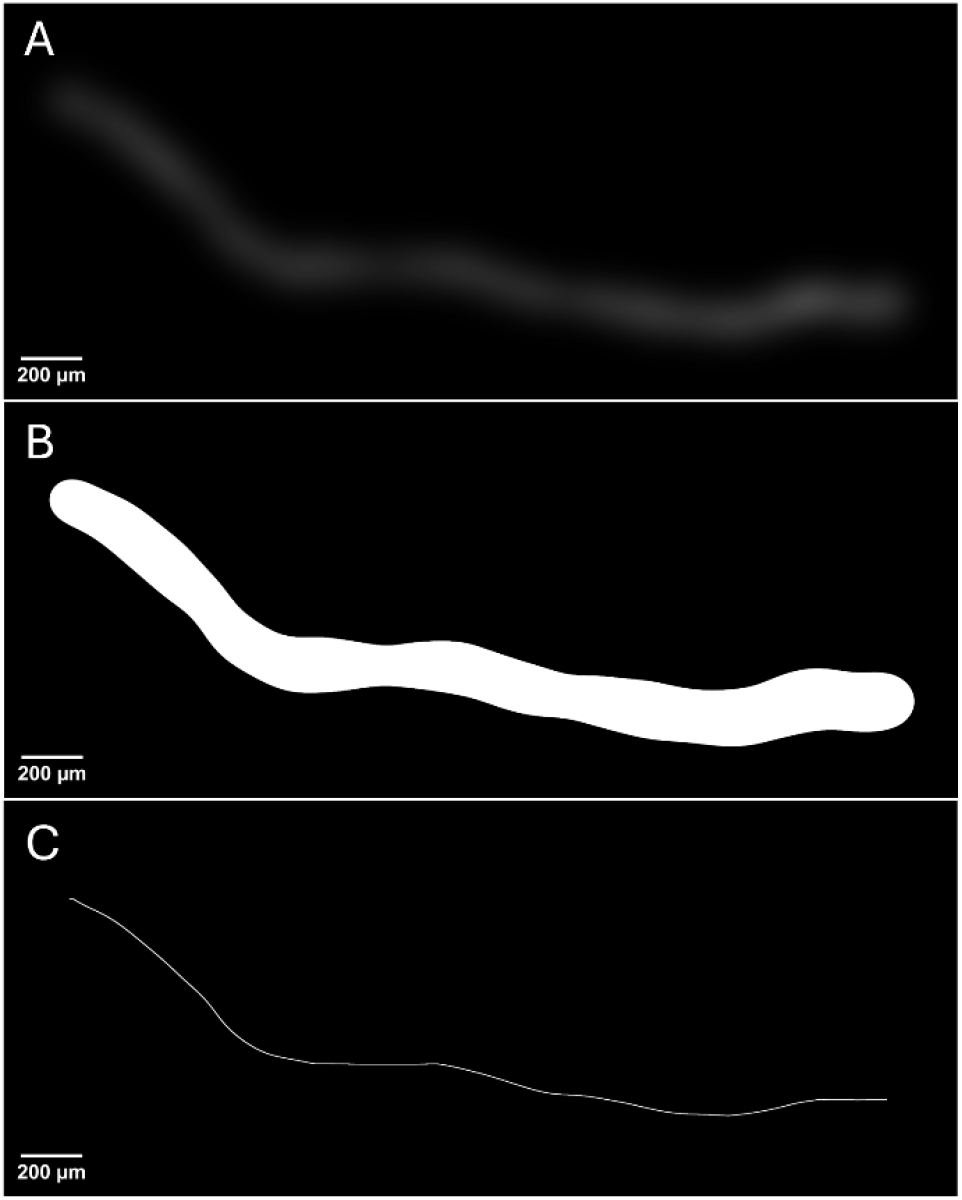
2D Fiber Skeleton Generation. **(A)** Gaussian-blurred nuclei mask (σ = 55 µm). **(B)** Gaussian-blurred nuclei mask following Otsu thresholding. **(C)** Skeleton for representation of fiber orientation (original skeleton width = 1 px; image has undergone 6 iterations of Dilation for visualization).

#### 4. File Structure and Organization

Outputs are organized automatically by subject, timepoint, and leg in nested directories.

### Python-based myonuclear analysis

We developed a Python based image analysis pipeline to quantify nuclear morphology, spatial position, orientation, and clustering within individual skeletal muscle fibers. The project was initially conceived as a hybrid *CellProfiler*–Python workflow, but the final implementation achieved greater flexibility and analytical control through a fully Python based design. The workflow processes each fiber’s input images—binary nuclear masks (STDIP), corresponding fiber skeletons, and z stacked fluorescence images—to extract per nucleus morphometric and spatial features.

The pipeline was designed to (1) ensure reproducible per-fiber processing, (2) maintain compatibility with existing file-naming and organizational conventions used in the Dreyer Lab, (3) enable flexible parameter tuning for microscope pixel scaling, depth spacing, and noise filtering, and (4) generate transparent outputs suitable for quality control and further statistical aggregation.

#### 1. Input structure and parsing

Each fiber is represented by three co-registered images stored within standardized subdirectories:

- *STDIP/* - binary standard-deviation projection image marking nuclear masks;
- *Skel/* - 2D representative fiber skeleton image;
- *TIFs/* - z-stacked 3D image with dimensions (Z, Y, X).

The parser is robust to naming variants and missing files.

#### 2. Parameterization

All adjustable constants—such as pixel-to-micron conversion, z-axis scaling, area filtering limits, and clustering radius—are stored in a *Params* dataclass. This allows all analysis steps to reference a single structured parameter object and supports command-line overrides; encapsulating parameters in a dataclass makes the pipeline reproducible and portable across imaging conditions and magnifications.

#### 3. Image loading and preprocessing

Utility functions handle file input and preparation:

- *_imread()* loads TIF/TIFF files using scikit-image.
- *_to_bool()* ensures binary STDIP masks are converted to Boolean arrays.
- *_label_mask()* labels nuclei in the binary image, optionally removing objects smaller than a given minimum area.

#### 4. Nuclear morphology measurements

The function *measure_nuclei_from_binary()* computes morphometric descriptors for each labeled nucleus of the STDIP image: area, perimeter, centroid coordinates, major/minor axis lengths, and orientation. Derived quantities include:

- *Aspect ratio* = minor/major axis length. A value of 1 indicates a perfect spheroid; values approaching 0 indicate an ellipsoid shape.
- *Shape class* (spherical / intermediate / ellipsoid) based on user-defined aspect-ratio thresholds.
- *Orientation* (deg) on 0–180° scale.

All measurements are then converted from pixel to micrometer units, specific to imaging protocol.

Nuclei outside user-defined area thresholds can be flagged and excluded to eliminate segmentation artifacts or merged/overlapping objects. Both included and excluded nuclei are retained for output in separate CSV files, allowing full traceability of excluded object.

Annotated mask overlays are also generated for quality control.

#### 5. Z-position assignment and Z-consistency filtering

Because widefield imaging introduced substantial Z-axis elongation artifacts, Z dimensions from microscope metadata could not be used for reliable 3-dimensional measurements or spatial clustering. To mitigate this distortion, the function *compute_mean_z_for_labels()* estimates the effective z-position of each nucleus directly from the z-stack intensity data. For each pixel coordinate in the stack, the index of maximal fluorescence intensity (z-argmax) is identified, producing a per-pixel map of maximal signal depth. For each labeled nucleus, the set of z-indices corresponding to its pixels is collected, and the mean z-position is accepted only when the within-object standard deviation of these indices is below a user-defined threshold (default = 2).

Additionally, to distinguish planar nuclei from vertically overlapping artifacts, the pipeline now applies a Z-consistency filter:

- *Included*: nuclei with low within-object Z standard deviation (planar; allows physiologic rouleaux)
- *Excluded*: nuclei with high Z-STD (vertical overlap / projection artifacts)

This yields a robust *Mean_Z_um* measure for all included nuclei and improves downstream 3D spatial analysis.

#### 6. Relative myonuclear orientation

While the preceding morphometric measurements (Section 4) describe nuclear orientation relative to the global x-y image axes, this does not capture the true alignment of nuclei with respect to individual muscle fibers, whose trajectories may curve or vary across the tissue section. To resolve this, the function *skeleton_angles_near_nuclei()* estimates the dominant local direction of each myofiber in the vicinity of every nucleus.

For each nuclear centroid, the algorithm identifies all skeleton pixels within a fixed radius and applies Principal Component Analysis (PCA) to their (x, y) coordinates. The first principal component corresponds to the dominant local axis of the fiber segment. Nuclear orientation, originally measured relative to the image x-axis, is then compared to this local fiber axis to compute a relative alignment angle (Figure 8).

**Figure 8:**
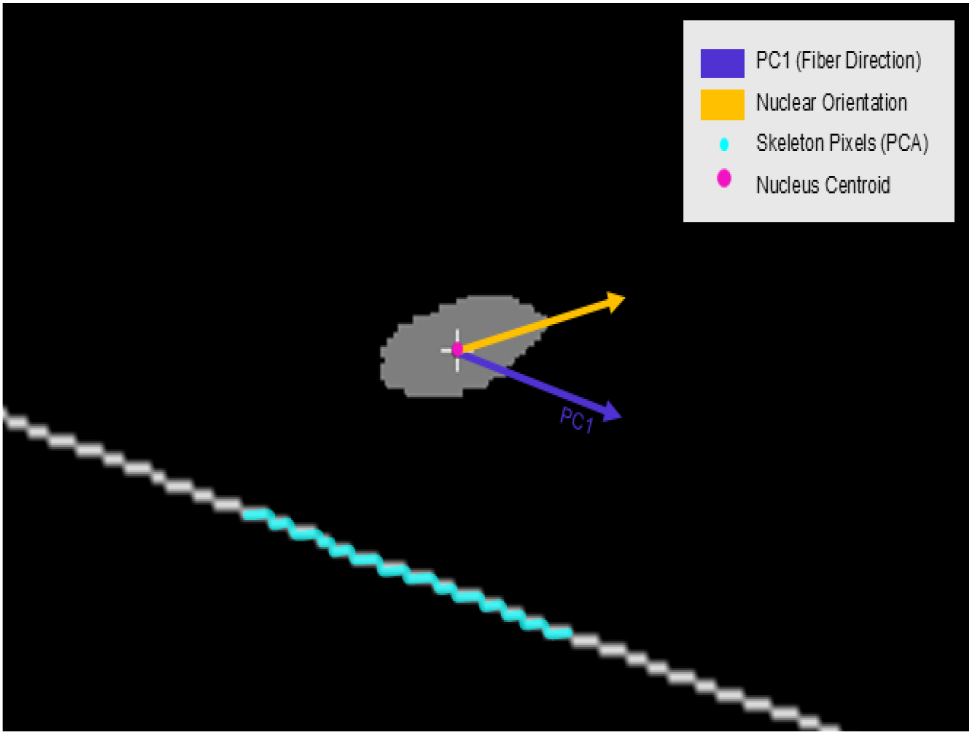
Raw Python generated image of principal component analysis (PCA) of local fiber direction. Local fiber direction is estimated by applying PCA to the coordinates of skeletonized fiber pixels within a fixed radius around each nucleus; the first principal component (PC1) indicates the dominant fiber direction.

This relative angle is expressed on a 0°–90° scale, where 0° denotes perfect alignment of the nucleus with the fiber axis and 90° indicates perpendicular orientation. This approach provides a curvature-aware measure of nuclear alignment that reflects local myofiber structure rather than global image geometry (Figure 9).

**Figure 9:**
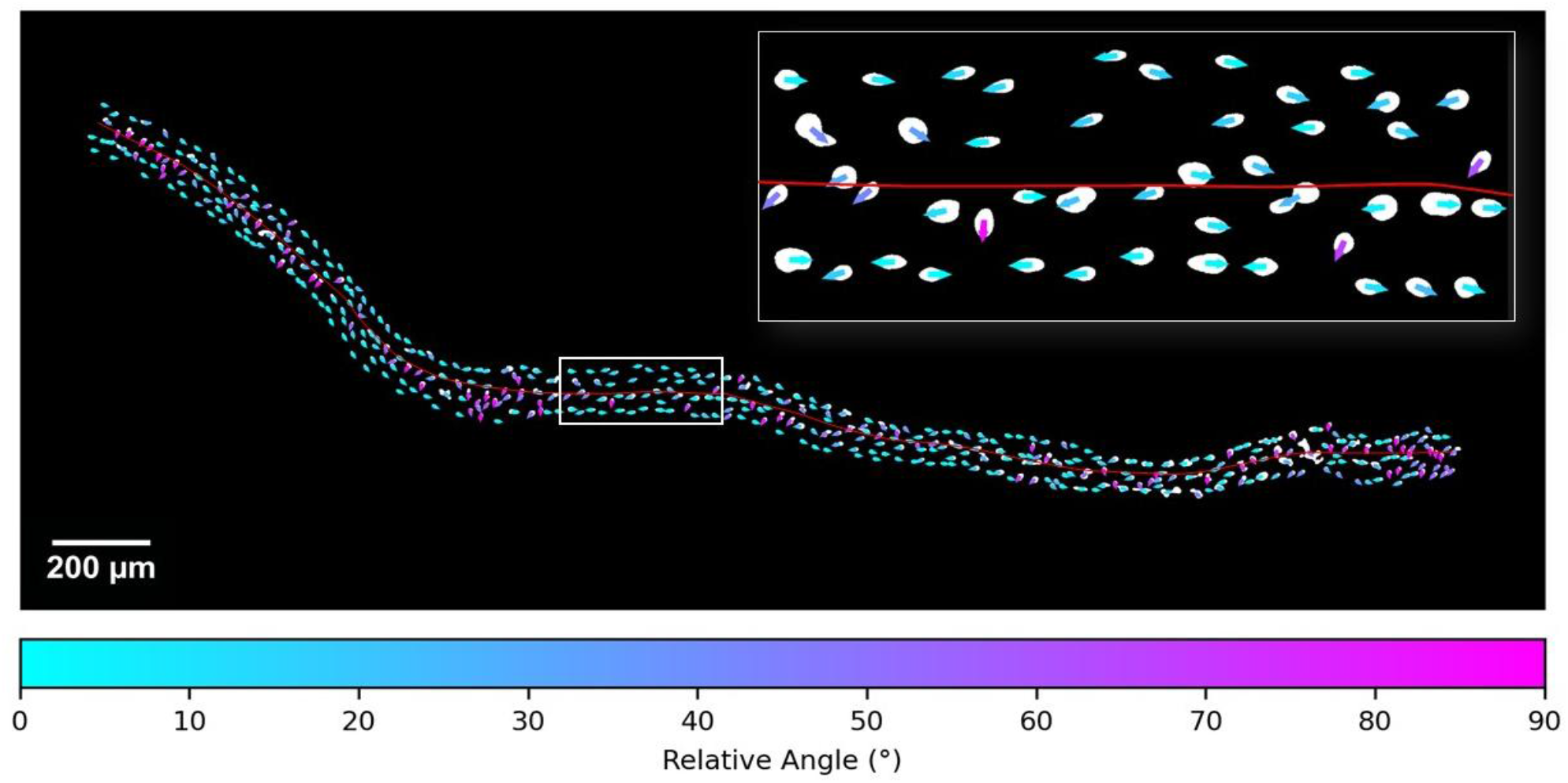
Raw Python generated image of relative orientation angles of myonuclei along a single myofiber. Each nucleus is overlayed with a colored arrow representing the orientation of the long axis of the nucleus to the long axis of the myofiber. A cyan arrow indicates parallel alignment, and a magenta arrow indicates perpendicular alignment to the long axis of the myofiber.

#### 7. Distance to fiber skeleton

To quantify nuclear positioning relative to the fiber core, *compute_distance_to_skeleton()* calculates the Euclidean distance from each nucleus centroid to the nearest skeleton pixel. This produces a *DistanceToSkel* variable (µm) for every nucleus. This metric enables downstream classification of “central” nuclei and supports rouleaux analysis.

#### 8. 3-dimensional clustering of nuclei

To identify spatial clusters of myonuclei within individual muscle fibers, we applied the Density-Based Spatial Clustering of Applications with Noise (DBSCAN) algorithm. DBSCAN was implemented using the scikit-learn library in Python. DBSCAN is an unsupervised density-based clustering algorithm that groups points with many nearby neighbors and marks outlier points that have few nearby neighbors. The clustering was performed on 3-dimensional myonuclear centroid coordinates (Position X, Position Y, Position Z).

The algorithm was parameterized with an *eps* value of 20 μm, which defines the maximum distance between two points for them to be considered part of the same neighborhood, and a *min_samples* value of 2, requiring at least three nuclei to form a valid cluster. These parameters were chosen based on the elbow method and observed spatial distribution of nuclei to ensure the detection of small but meaningful nuclear groupings.

Cluster labels assigned by DBSCAN were then added to the dataset. Nuclei not assigned to any cluster (classified as noise) were labeled as -1 by the algorithm and recorded accordingly (Figure 10).

**Figure 10:**
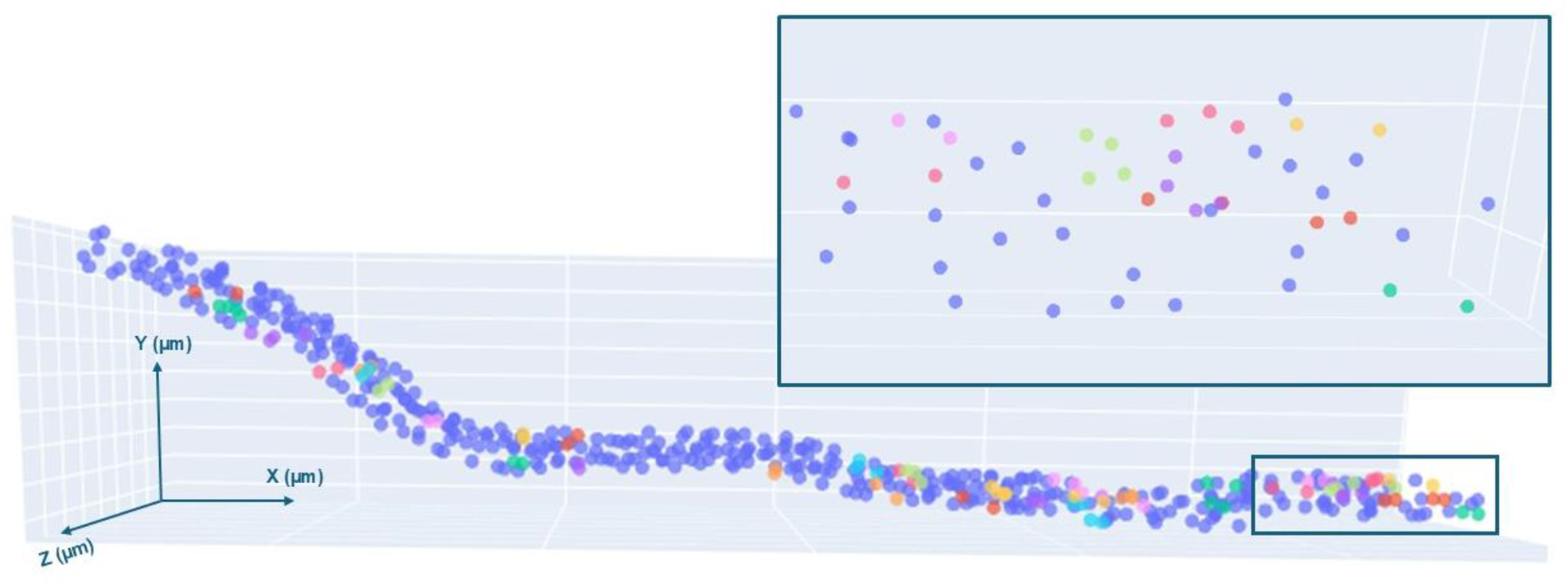
Raw Python generated image of 3D myonuclear spatial analysis with DBSCAN. Cluster labels assigned by DBSCAN visualized using Plotly in a scatter plot. Each myonucleus is colored according to its cluster identity; nuclei not assigned to any cluster are labeled as -1 and displayed in the plot in gray.

#### 9. Nearest-neighbor analysis

To further characterize the local nuclear microenvironment, the pipeline computes 3- and 5-nearest-neighbor (NN3/NN5) features for each included nucleus using 3D Euclidean distances. Extracted contextual features include:

- *NNk_MeanDist_um* – local nuclear packing density
- *NNk_MeanAspectRatio* – neighborhood shape profile
- *NNk_Frac_Spherical / Ellipsoid / Intermediate* – shape class composition
- *NNk_Frac_Parallel* / Perpendicular / IntermediateOrientation – orientation coherence within the microenvironment
- *NNk_MeanRelAngle_deg* – neighbor alignment relative to local fiber axis

These metrics quantify nuclear neighborhood structure and support analysis of “myonuclear footprint” organization.

#### 10. Fiber diameter profiling

The function *compute_fiber_width_profile()* samples diameters along the skeleton at fixed intervals (default 100 µm). Diameters are estimated by projecting included nuclei positions perpendicular to the local axis and measuring left– right separation. This yields a diameter profile and summary statistics (mean, min, max, SD) to inform downstream volume analyses.

#### 11. Myonuclear domain volume

The pipeline also computes a volume-based myonuclear domain for each fiber:

- *FiberVolume* (µm^3^) = π·(MeanDiameter/2)^2^ · ImarisFiberLength
- *DomainVolume_um3_Python* = FiberVolume / PythonMyonuclei
- *DomainVolume_um3_Imaris* = FiberVolume / ImarisMyonuclei (if available)

This provides a volumetric estimate of the cytoplasmic territory associated with each myonucleus.

#### 12. Biopsy-level summaries

For each fiber, derived results are merged with *Imaris* master file metrics (fiber length, nuclei counts), if available. Additional biopsy-leve variables are computed:

- Nuclear counts (PythonMyonuclei, ImarisMyonuclei)
- Diameter statistics and FiberVolume
- Myonuclear domain volume (Python/*Imaris*)
- Neighbor microenvironment metrics (fiber-level means of NN3 features)
- Shape and orientation distributions
- Cluster# (3D DBSCAN clusters)
- Rouleaux and central-rouleaux counts

#### 13. Batch orchestration and outputs

Helper functions automate batch processing across subjects, timepoints, and legs. Each subject receives a workbook *(<Subject>_results*.*xlsx)* with one sheet per timepoint, preserving column order and preventing Excel auto formatting of fiber IDs. Per fiber outputs include:

- *nuclei_results*.*csv* (included nuclei)
- *excluded_nuclei*.*csv* (excluded objects)
- *fiber_width_profile*.*csv* (diameter sampling)
- *overlay*.*png* (annotated mask image)

Biopsy level summaries integrate Python metrics with outputs, providing a unified dataset for downstream statistical analysis.

This workflow integrates morphometric, orientational, and 3D spatial analyses of single-fiber nuclei into a unified Python framework. By combining region-property extraction, Z-position inference, curvature-aware orientation estimation, diameter profiling, nearest-neighbor microenvironment analysis, and unsupervised clustering, the pipeline yields a rich quantitative description of myonuclear organization without the need for proprietary software. The modular design enables adaptation to a range of staining protocols and imaging systems while producing fully auditable outputs for downstream statistical modeling and biological interpretation.

## Discussion

In this study, we developed and validated a semi-automated pipeline analysis method to quantify myonuclei number and structural features along the length of individual single muscle fibers from biopsy. Our approach allows for reproducible, high-resolution measurement of myonuclear number, morphology, clustering, and orientation from immunohistochemical images. By integrating myonuclear density metrics with spatial orientation of similarly shaped myonuclei, our method provides a foundation for examining how nuclear organization relates to multinucleated muscle cell plasticity.

Although several automated pipelines exist for analyzing muscle ultrastructure, most are designed for tissue cross-sections, where sampling many fibers simultaneously increases statistical confidence at the tissue level. In contrast, our motivation was to maximize individual muscle cell-specific power to quantify myonuclear accretion and local and longitudinal organization. This required sampling along the length of single fibers to capture the true spatial relationships of nuclei that cannot be inferred from cross-sectional preparations.

Our first aim was to achieve accurate quantification of myonuclear number along the length of individual single fibers, enabling precise density metrics such as myonuclei per millimeter and myonuclei per volume. By combining FIJI-based preprocessing with Python analysis and integrating by integrating fiber-length measurements from *Imaris*, our pipeline generates reproducible nuclear counts that ensures reproducible quantifiable data. This approach provides confidence in density estimates that are difficult to achieve with manual methods. Compared to cross-sectional imaging pipelines, which maximize statistical power by sampling many fibers simultaneously, our longitudinal approach prioritizes fidelity in capturing nuclear accretion and local organization along the fiber axis. Limitations include reliance on widefield microscopy, which prevents true 3D morphology assessment, and the need for manual imaging, which remains time-intensive and costly. Nonetheless, the reproducibility and accuracy of density metrics represent a significant advance over manual quantitation.

A second aim was to develop a scalable method for representing merged DAPI z-stacks into a standardized format suited for batch-processing. Because widefield z-stacks generate large, computationally demanding datasets, our FIJI macro reduces file size by generating standardized 2D projections while preserving nuclear structural information. This design allows efficient downstream analysis and maintains compatibility with existing file-naming conventions. While some 3D detail is lost and occasional nuclear overlap can lead to underestimation, our macro offers a practical balance between fidelity, scalability, and computational efficiency—making longitudinal single-fiber nuclear analysis feasible across large cohorts. Limitations include occasional overlap of nuclei that cannot be fully separated by watershed segmentation, leading to underestimation of nuclear counts.

The third aim was to establish a reproducible framework for quantifying nuclear morphology, orientation, clustering, and spatial relationships from 2D z-projections. The Python implementation extracts morphometric parameters from binary nuclear masks—including area, perimeter, centroid location, axis lengths, aspect ratio, and shape class—producing standardized metrics for comparisons across fibers or conditions. Orientation is calculated relative to the local fiber axis, determined via principal component analysis of nearby skeleton pixels that define trajectory with nuclear orientation, yielding alignment angles (0– 90°) that account for fiber curvature and not global image axes.

Spatial organization is characterized through DBSCAN clustering of three-dimensional centroid coordinates, which identifies discrete nuclear clusters and labels noise nuclei, and through centroid-to-skeleton distances that enable classification of central nuclei and central Rouleaux formations. Fiber organization is further interrogated with Nearest-Neighbor statistics, which utilizes local density patterns to reveal whether the nuclei tend towards random, clustered, or ordered distributions.

Together, these measures generate a reproducible “fingerprint” for each muscle fiber, integrating morphometric, orientational, and spatial descriptors into a unified profile. This fingerprinting framework allows for longitudinal and comparative studies, where distinct nuclear arrangements such as linear rouleaux formations of nuclear chains may be correlated with biological variables including age, disuse, or exercise status. Compared with existing tools—such as *CellProfiler*, which provides accessible workflows for segmentation and basic morphometry, *Imaris* offers highly accurate volumetric analyses within a proprietary environment. Our pipeline complements these approaches by emphasizing transparency, reproducibility, and scalability for widefield datasets. Although widefield imaging and occasional segmentation inaccuracies remain limitations, the approach provides systematic and high-throughput quantification of nuclear organization in two dimensions.

In conclusion, we present a semi-automated pipeline that (1) quantifies myonuclear number along single muscle fibers; (2) classifies nuclei into morphological categories (spherical, intermediate, elongated); (3) quantifies the number of similarly shaped nuclei within a defined spatial radius; and (4) determines nuclear orientation relative to the local fiber axis. To the best of our knowledge, this is the first integrated pipeline to apply widefield microscopy—common in most laboratories—to longitudinal single-fiber nuclear analysis. Designed to process large numbers of composite images, this method yields rich datasets describing myonuclear distribution, morphology, and spatial relationships along the fiber. This method has potential for broad application in muscle biology and other fields requiring quantitative analysis of subcellular spatial organization.

## Supporting information

Dreyer_SF Pipeline Supplemental

## Acknowledgements

We would like to acknowledge the Genomics & Cell Characterization Core Facility at the University of Oregon. We thank members of the Dreyer Lab for comments and suggestions while testing our pipeline and drafting this manuscript.

## Data and materials availability

All data and materials are available in the manuscript and supplemental protocol. Code is available at: https://github.com/DreyerLabUO/23Wu-SingleFiber.

