## Supplementary material for "A semi-automated pipeline for morphological analysis of myonuclei along single muscle fibers": Dreyer_SF Pipeline Supplemental

#### Table of Contents


I. **3D Single Fiber Myonuclei & Length Quantification with *Imaris***
   a. **Pre-Processing**
   b. **Myonuclear Quantification (*Spots*)**
   c. **Fiber Segmentation for Length Measurements (*Surfaces*)**
   d. **Fiber Length Measurement (*Filaments*)**
II. **2D Image Processing with *FIJI Macro***
   a. **Macro Import and Set-Up**
   b. **Raw TIF Export**
   c. **Z-Projection & Nuclei Mask Generation**
   d. **Fiber Skeleton Generation**
III. **Analysis with *Python***
   a. **Running the script**
   b. **Hierarchical *.csv* Output**
IV. **Our Computer Specifications**

### I. 3D Single Fiber Myonuclei and Fiber Length Quantification using *Imaris*

#### A) *Z-Stack Import and Processing*

**Procedure:**

1. Convert raw Leica .lif files into Imaris .ims format using the Imaris File Converter.
2. Import merged z-stacks (files ending in "_Merged") to ensure full fiber coverage.
3. Adjust DAPI channel display (optional: change to cyan for improved contrast).
4. Apply background subtraction with a filter width of 10 μm.
5. Set intensity threshold at the histogram "elbow" to suppress background illumination.

**Rationale:** Imaris requires .ims file types to load images. Z-stacks acquired with a Leica Thunder widefield microscope are saved as .lif files and must be converted to .ims format to enable downstream analysis. These preprocessing steps are necessary to enhance the DAPI signal relative to the background. Background subtraction eliminates significant background illumination, and thresholding improves the DAPI signal appearance for increased ease of distinguishing myonuclei from background by eye.

**Adaptations:**

- Adjust the filter width to the average diameter of the objects of interest (nuclei).
- For datasets with stronger background illumination, raise the intensity threshold. For cleaner datasets, thresholding may be relaxed to preserve faint nuclear signals.

#### B) *Myonuclear Quantification using Imaris' Spots module*

**Procedure:**

1. Open the *Spots* tool in Imaris.
2. Disable default detection settings.
3. Set *Estimated XY Diameter* to 10 μm.
4. Adjust *Quality Threshold* between background and nuclear peaks.
5. Record Spot count as *Imaris Myonuclei*.
6. Perform manual Quality Control (QC): note missed nuclei (false negatives, *Missed Myonuclei*) and artificial *Spots* (false positives, *Artificial Myonuclei*).

**Rationale:** Spot detection in Imaris is semi-automated. The software estimates myonuclei based on signal contrast and size, but manual verification is required to correct for missed or artificial identifications. Missed or artificial myonuclei are more common in low-quality, noisy images.

**Adaptations:**

- Adjust *Estimated XY Diameter* to fit the average diameter of your object of interest if it differs greatly from 10 µm.
- This process can be fully automated with the use of *Imaris' Pixel Classifier,* which allows you to train a proprietary model for your data set that can offer greater results compared to the standard *Quality Theshold* with Otsu.
    - We chose to stick with Manual Quality Control over the *Pixel Classifier* to preserve the highest standard of myonuclear quantification, but the *Pixel Classifier* is a great option for those without the available time to manually double-check each dataset.

#### C) *Fiber Segmentation using Imaris' Surfaces module*

**Procedure:**

1. Open the *Surfaces* tool in Imaris.
2. Disable default settings.
3. Smooth the DAPI channel surface with detail width 12 µm.
4. Apply absolute intensity thresholding until fiber is encompassed.
5. Mask the DAPI channel: set voxel intensity inside surface to 1.00, outside to 0.00.

**Rationale:** Imaris' Filaments module requires consistent signal to generate a smooth line, which this method uses to accurately measure fiber length. This step essentially creates a binary mask of the fiber in 3D, in turn generating a consistent signal throughout the fiber that enables Filaments to produce accurate fiber length measurements.

**Adaptations:**

- Adjust smoothing width for thinner or thicker fibers.
- Fine-tune thresholding for uneven illumination.
- Increase threshold if masking produces incomplete coverage.

#### D) *Fiber Length Measurement using Imaris' Filaments module*

**Procedure:**

1. Open the *Filaments* tool in Imaris.
2. Select "No Loops No Soma" detection type.

3. Use masked DAPI channel as input.
4. Indicate thinnest fiber diameter as 100 µm.
5. Switch to Slicer View (slice width = 10 µm).
6. Place seed points along fiber centerline end-to-end.
7. Generate Filament.
8. If Filament does not capture curvature precisely, go back to Seed Points (Step 3/6 in Filaments module) and adjust placement until Filament adequately represents fiber orientation.
9. Once Filament is adequate, record Filament length in microns (found under Statistics tab within *Filaments* module).

**Rationale:** Imaris' Filaments module requires consistent signal to generate a smooth line, which this method uses to accurately measure fiber length. By placing representative seed points along the fiber center, the filament path follows the true centerline of the fiber, ensuring accurate length measurement. When placing Seed Points, slice diameter is set to 10 microns to ensure that the center of the Seed Points (and subsequent Filament) do not deviate more than 10 microns from the center of the fiber.

**Adaptations:**

- Adjust thinnest diameter for smaller or larger fibers.
- Increase seed density for curved fibers. Fewer seed points suffice for shorter fibers.

#### II. Automated 2D Image Processing using *FIJI* Macro

##### A) *Macro Import and Set-Up*

**Procedure:**

1. Download Fiji/ImageJ and enable Bio-Formats update site.
2. Launch Fiji and import the macro (.*ijm*) file.
3. Create an input folder to contain your project (.*lif*) files, and create an output folder for processed images to be saved to.
4. Click "Run" on the macro & select input/output directories when prompted.
5. Once macro finishes, double-check the output images for abnormal processing results.

**Rationale:** Metadata parsing organizes outputs systematically. Restricting to merged stacks ensures only the merged images containing fiber segments of interest are imaged, and that individual tiles (not TileScan) images are skipped.

**Adaptations:** Our image stacks of interest (merged TileScan z-stacks acquired with Leica equipment) are named using a standardized naming convention to allow the macro to recognize the file and save it to the proper location. To adjust the macro to read your files, please adjust the Regex pattern to match your file naming structure. In our Fiji macro, lines __ - __ are responsible for this file parsing and nested directory creation.

Standard Regex Pattern: [StudyCode]_[SubjectID]_[Timepoint]_[Leg]_Merged.*lif*

Example Regex Pattern: 23_Wu_01_D14_L_Merged.lif

The Regex pattern above is for a biopsy obtained for our 23_Wu study, from Subject 01, at our D14 timepoint, from the Subject's left leg (L).

##### B) *Raw TIF File Saving*

**Procedure: *No user input required, completed automatically by macro***

**Rationale:** A raw TIF of each stack analyzed by the macro is saved prior to Z-projection and image processing. This raw TIF is used downstream to identify the z-position of myonuclei masks on the 2D projection, utilizing the XY positions of the masks to locate nuclei in the raw TIF and save their respective Z-locations.

**Adaptations:**

- This step can be skipped if 3D nuclei orientation and colocalization is not necessary for your analysis.

##### C) *STD Z-Projection and Nuclei Mask Generation*

**Procedure: *No user input required, macro steps detailed below***

1. Apply rolling ball background subtraction (radius = 30 px).
2. Collapse z-stack into 2D standard deviation projection (STDIP).
3. Apply median filter (radius = 1 px).
4. Threshold using Otsu's method.
5. Binarize and save mask in "STDIP" directory.

**Rationale:** Background subtraction with a rolling ball radius of 30 pixels, our estimated average diameter of a myonucleus, is applied to the entire z-stack to remove low-frequency background signal. The Standard Deviation Z-projection emphasizes high-intensity regions in a single 2D plane, allowing for 2D representation of all myonuclei. A median filter further suppresses noise, and Otsu thresholding provides reproducible segmentation by isolating the high-intensity objects from a low-intensity background to create a binary 8-bit mask of the nuclei.

**Adaptations:**

- Adjust rolling ball radius for different nuclear sizes.
- Increase median filter radius for noisier datasets.
- Use alternative thresholding methods if Otsu fails to separate signal.

##### D) *STD Z-Projection and Nuclei Mask Generation*

**Procedure: *No user input required, macro steps detailed below***

1. Duplicate STDIP mask.
2. Apply Gaussian blur (σ = 55 μm).
3. Threshold with Otsu's method.
4. Isolate large objects (>70,000 $px^2$).
5. Skeletonize fiber mask and save in "Skel" directory.

**Rationale:** This step transforms the DAPI mask into a representative mask of the overall fiber shape. Large objects are isolated to remove smaller artifacts, and the resulting binary mask is skeletonized to represent fiber orientation and length.

**Adaptations:**

- Adjust Gaussian blur filter width for thicker or thinner fibers if needed; thin fibers require a smaller sigma value and thicker fibers require a larger value.

#### III. Analysis with *Python*

##### A) *Run the Python Analysis*

**Procedure:**

See GitHub repository for additional documentation: https://github.com/DreyerLabUO/23Wu-SingleFiber.

1. Open a terminal (Anaconda Prompt recommended).
2. Create and activate a Conda environment:

```
conda create --name sfpipeline python=3.10
conda activate sfpipeline
```

3. Install the required Python packages/dependencies:

```
python -m pip install numpy pandas scikit-image scikit-learn openpyxl Pillow
```

4. Save the script SF_analysis_pipeline.py in your working directory.
5. Run the pipeline.

   From the activated Conda environment terminal, run the script with desired parameters:

```
python SF_analysis_pipeline.py "D:\MacroOutput" --imaris_master
"C:\path\to\Imaris_master.xlsx" \
--pixel_size_xy_um 0.329 \
--z_scale_um_per_index 2.7 \
--min_area_px 200 \
--max_area_px 0 \
--z_std_threshold 2.0 \
--dbscan_eps_um 20 \
--skeleton_radius_px 20 \
--fiber_width_step_um 100 \
--fiber_width_max_radius_um 100
```

   The base directory (e.g., D:\MacroOutput) must contain subject folders, each with timepoints, legs, and outputs from FIJI:

```
Subject/
   Timepoint/
         L or R/
               SDTIP/
               Skel/
               TIFs/
```

**Rationale:** Once the FIJI/ImageJ macro has been executed and the SDTIP masks, skeleton images, and z-stacks are organized into the standard directory structure, the entire single-fiber analysis can be executed using the Python pipeline.

**Adaptation:**

***Recommended User-Adjustable Parameters***

| Parameter | Default | What It Controls | When to Adjust | Typical Range / Guidance |
|---|---|---|---|---|
| --pixel_size_xy_um | 1.0 | Microns per pixel in XY plane | Always adjust to match microscope objective and camera | 0.05–0.65 µm depending on magnification |
| --z_scale_um_per_index | 2.7 | Z-step spacing (µm per slice) | Always adjust based on acquisition z-step | Usually 1–5 µm |
| --min_area_px | 0 | Minimum nuclear area ($px^2$) for inclusion | Increase if debris/small fragments are detected | Often 100–400 $px^2$ |
| --max_area_px | 0 | Maximum nuclear area ($px^2$) for inclusion | Set if large merged nuclei appear in segmentation | 800–2000 $px^2$ depending on magnification |
| --z_std_threshold | 2.0 | Z-consistency filter; excludes vertically elongated or overlapping nuclei | Tighten if z-stacks are noisy; loosen if intended nuclei are excluded | 1.5–3.0 typical |
| --dbscan_eps_um | 20 | 3D distance threshold for DBSCAN clustering | Change if clusters are too fragmented or too large | 10–30 µm |
| --dbscan_min_samples | 2 | Minimum nuclei to form a cluster | Increase to require more robust clusters | 2–4 typical |
| --skeleton_radius_px | 20 | Neighborhood around each nucleus used for PCA-based local axis estimation | Increase if skeleton is thin or broken; decrease if fibers curve sharply | 10–40 px |
| --fiber_width_step_um | 100 | Interval (µm) for sampling diameter along the fiber | Reduce if high-resolution diameter profile is desired | 50–200 µm |
| --fiber_width_max_radius_um | 100 | Maximum search radius for estimating fiber boundaries using nuclei positions | Increase for thick fibers; decrease if nuclei are sparse | 50–150 µm |
| --dbscan_eps_um | 20 | Distance threshold for clustering | Adjust based on nuclear packing density | 10–30 µm |
| *(internal behavior)* | — | NN3/NN5 nearest-neighbor analysis | Auto; no tuning needed unless downsampling nuclei | Not user-adjusted |

#### ***B) Output Files and Their Interpretation***

The pipeline produces structured outputs at two levels: per-fiber and per-subject.

**Per-Fiber Outputs**

Located in directories of the form: <Subject>/<Timepoint>/<Side>/<FiberTag>_output/

1. nuclei_results.csv. Contains all included nuclei, with:

- Morphometrics
- Z-consistency metrics
- Distance to skeleton
- Orientation relative to fiber axis
- 3D DBSCAN cluster labels
- Nearest-neighbor microenvironment features (NN3 & NN5)
- Shape and orientation context of neighboring nuclei

2. excluded_nuclei.csv. Lists nuclei removed due to:
    - Area thresholds
    - Excessive Z-STD ("vertical overlap")
    - Other exclusion criteria

   Includes explicit exclusion reasons.
3. fiber_width_profile.csv. Diameter measurements sampled along the fiber length, using included nuclei.
4. overlay.png. Visualization of all labeled nuclei on the SDTIP mask.

**Per-Leg Biopsy Summary**

The script generates a CSV: <Subject>_<Timepoint>_<Side>_biopsy_summary.csv
Containing per-fiber summary metrics:

- Python and Imaris myonuclei counts
- Diameter statistics
- FiberVolume (cylindrical approximation)
- Myonuclear domain volume (Python and *Imaris*-based)
- Fiber-level means of NN3 neighbor features
- Shape and orientation distributions
- Rouleaux and central-rouleaux counts

Unmatched fibers (no Imaris entry) are written to a companion: *_excluded_fibers.csv.

**Subject Workbook**

Each subject receives <Subject>_results.xlsx with one sheet per timepoint, containing all per-nucleus features, including NN metrics, Z positions, cluster IDs, and orientation data.

### IV. Our Computer Specifications

This pipeline was trained and developed on both a standard, in-lab desktop workstation and a high-performance computer operated in the Genomics and Cell Characterization Core Facility (G3CF) at the University of Oregon.

> ***Lab desktop*: Dell Precision 3630 Tower workstation running Microsoft Windows 11 Enterprise (Build 26100). The system is equipped with an Intel® Core™ i7-9700 CPU @ 3.00 GHz with 8 cores and 8 threads and operates in UEFI mode with Secure Boot enabled. The machine has a 64-bit architecture and BIOS version 2.26.0 (dated December 8, 2023).

> ***G3CF computer*: 32-core AMD Ryzen Threadripper 3970X processor running at 3.9 GHz and 256 GB of DDR4 RAM. The system features a custom water-cooled configuration optimized for high-throughput image processing and is powered by an NVIDIA GeForce RTX 5090 graphics card to support GPU-accelerated rendering and analysis tasks. The machine runs Microsoft Windows 10 Enterprise (version 10.0.19045, build 19045) in UEFI BIOS mode. The software environment includes Imaris 9.9 with the Cell and Filaments Modules Imaris 9.9 Converter, MATLAB 2018b, and Zeiss Zen Black.
